# B cells suppress medullary granulopoiesis by an extracellular glycosylation-dependent mechanism

**DOI:** 10.1101/604264

**Authors:** Eric E. Irons, Melissa M. Lee-Sundlov, Karin M. Hoffmeister, Joseph T.Y. Lau

## Abstract

The immune response relies on the timely integration of cell-intrinsic processes with cell-extrinsic cues. During infection, B cells vacate the bone marrow for the emergency generation of granulocytes. However, it is unclear if cross-talk between B cells and neutrophils also encourages the return to homeostasis. Here, we report that B cells remodel glycans on hematopoietic progenitors to suppress granulopoiesis. Human B cells secrete active ST6Gal-1 sialyltransferase to modify the sialylation and Gr-1 expression of co-cultured hematopoietic progenitors. After adoptive transfer, total hematopoietic and B cells modified the sialylation of non-self cells and elevated blood ST6Gal-1. Mature IgD+ B cells co-localized with megakaryocytes to sialylated bone marrow niches, suggesting their role in medullary extrinsic sialylation. Finally, ST6Gal-1 expression in multiple myeloma cells negatively correlated with neutrophil abundance in human patients. Our results highlight the growing significance of extracellular glycoslytransferases as mediators of a novel glycan-dependent interaction between B cells and granulocytes.

## Introduction

Hematopoiesis generates the blood cells necessary for gas exchange, hemostasis, and immune defense. Cell-intrinsic developmental programs orchestrate these lineage decisions, but they are guided by systemic signals to convey the dynamically changing needs for specific cell types [1]. Dysregulated communication of such extrinsic cues results in imbalanced blood cell production and can trigger pathologic processes including anemia, thrombocytopenia, inflammation, and autoimmunity [2; 3]. In malignancy, the disruption of normal differentiation within hematopoietic stem and progenitor cells by tumors can lead to insufficiencies in one or more blood cell lineages, a common complication [4].

The sialyltransferase ST6Gal-1 is a glycan-modifying enzyme mediating the attachment of α2,6-sialic acids. Canonically, it resides within the intracellular ER-Golgi secretory apparatus, but there is also an extracellular blood-borne form [5; 6; 7]. In addition to ST6Gal-1, a number of other terminal glycosyltransfearases are also present in systemic circulation [8]. Fluctuating levels of blood-borne ST6Gal-1 are associated with a wide array of conditions, especially metastatic cancers, where the level of blood-borne enzyme is often associated with poorer patient outcomes [9; 10; 11; 12; 13; 14; 15; 16]. Early reports also associated elevated blood ST6Gal-1 with systemic inflammation [17; 18], atherosclerosis [19], Alzheimer’s disease [20], and alcohol-induced liver disease [21; 22; 23; 24]. The physiologic contributions of extracellular ST6Gal-1 in these diseases, however, remain poorly understood. We have hypothesized that secreted ST6Gal-1 can access distant sites to modify circulating plasma components and surfaces of target cells that do not express ST6Gal-1 [25; 26]. Previously, we observed that blood-borne ST6Gal-1 can profoundly modify leukocyte differentiation by attenuating G-CSF dependent granulocyte production [27] while promoting BAFF-dependent survival in B cells [28]. In mouse models, circulatory ST6Gal-1 insufficiency results in an exuberant granulocytic inflammatory response [29; 30] that can be therapeutically ameliorated by intravenous infusion of recombinant ST6Gal-1 [31]. Extracellular, liver-derived ST6Gal-1 is also a major determinant of serum immunoglobulin G sialylation, which activates anti-inflammatory pathways within innate immune cells through Fc receptors [32; 33; 34]. In the periphery, activated platelets supply the necessary sugar donor-substrate to support such extrinsic glycosylation reactions [8; 26; 35; 36], but the source of extracellular sugar donor-substrate within the marrow medullary spaces is not known.

The principal source of extracellular ST6Gal-1 in circulation is believed to be the liver [18; 37; 38], where ST6Gal-1 expression is activated by glucocorticoids and IL-6 [18; 39; 40; 41]. However, B cells also robustly express ST6Gal-1, which synthesizes the ligands for sialic acid-binding receptors CD22 and Siglec-G [42; 43]. Here, we report that hematopoietic lineage cells, particularly B cells, contribute significantly to the extracellular pool of functional ST6Gal-1 and the sialylation of non-self cells both *in vitro* and *in vivo*. B cells secrete functionally active ST6Gal-1 that sialylates hematopoietic progenitors in co-culture and suppresses granulopoietic differentiation. *In vivo*, we observed consistent and frequent co-localization of IgD+ mature B cells and PF4+ megakaryocytes in richly α2,6-sialylated niches of the bone marrow. In bone marrow specimens of treatment-naïve human multiple myeloma, we observed a striking negative association between marrow plasma cell ST6Gal-1 expression and prevalence of bone marrow neutrophils. Our study is the first to demonstrate that the liver is not the sole source of ST6Gal-1 responsible for extracellular sialylation, and underscores a novel relationship between B lymphocytes and hematopoietic progenitors influencing neutrophil production in the marrow.

## Results

### Human B lymphoblastoid cells secrete enzymatically active ST6Gal-1

B cell expression of ST6Gal-1 is critical for B cell development and function secondary to engagement of the lineage-specific lectin CD22 with α2,6-sialic acid [44]. Although ST6Gal-1 is expressed in multiple tissue and cell types, it is thought that secreted, extracellular ST6Gal-1 is exclusively derived from the liver, as a hepatocyte conditional knockout of ST6Gal-1 results in vastly reduced serum ST6Gal-1 activity [37]. In addition to hepatocytes, ST6Gal-1 is expressed strongly in mature B cells [42], although numerous other cell types also express ST6Gal-1 to varying degrees [45]. The ability of non-hepatic cells to secrete functional ST6Gal-1 to drive extrinsic sialylation has not been formally studied.

We have recently analyzed the expression of ST6Gal-1 within bone marrow and splenic B cell populations in mice and found that maximal ST6Gal-1 expression occurred in transitional and mature stages of development [28]. In order to assess if human B cells are capable of secreting ST6Gal-1, we analyzed four B cell lines representing pre-B cell receptor (NALM-6), B cell receptor expressing (Louckes), and post-B cell receptor (RPMI 8226, MM1.S) developmental stages. ST6Gal-1 mRNA expression was detectable in all cell lines except RPMI 8226, with highest expression in the germinal center line. This is consistent with our previous observation that BCR activation induces ST6Gal-1 expression [46]. Expression of the β-site amyloid precursor protein-cleaving enzyme 1 (BACE1), believed to be required to liberate the secreted ST6Gal-1 domain from its N-terminal membrane anchor [47; 48], was detectable within all lines except the plasma cell MM1.S (*Fig 1a*). The pattern of ST6Gal-1 expression was confirmed on the intracellular level by western blot (*Fig 1b*). To assess the size and functional activity of secreted ST6Gal-1, B lymphoblastoid cell lines were seeded in serum-free medium for 3 days, and ST6Gal-1 in the conditioned medium analyzed by western blot and assayed for sialyltransferase activity towards an artificial acceptor. ST6Gal-1 protein was present in the conditioned medium of all expressing cell lines, and accumulated with time (*Fig 1c*). B cells expressing BACE1 secreted a 42-kDa, soluble form of ST6Gal-1 (NALM-6, Louckes), consistent with the proteolysis of the full-length protein. Unexpectedly, all cells also secreted the 50-kDa, full-length form of ST6Gal-1. This was the predominant secreted form in MM1.S cells, which express minimal levels of BACE1. In conditioned medium from day 3 of culture, α2,6-sialyltransferase activity generally agreed with protein analyses, whereas α2,3-sialyltransferase activity varied independently (*Fig 1d*). These results demonstrate that human B cell lines are capable of secreting ST6Gal-1 *in vitro*. Moreover, the secretion of the apparently full-length ST6Gal-1 form from MM1.S in the absence of BACE1 was also of interest, as the canonical mechanism of glycosyltransferase release from the intracellular Golgi membrane involves proteolysis of the exposed stem joining the catalytic domain with the N-terminal transmembrane anchor. Although to the best of our knowledge, the secretion of unprocessed, full-length ST6Gal-1 has never been reported, its potential biologic significance is not explored further here.

**Figure 1.**
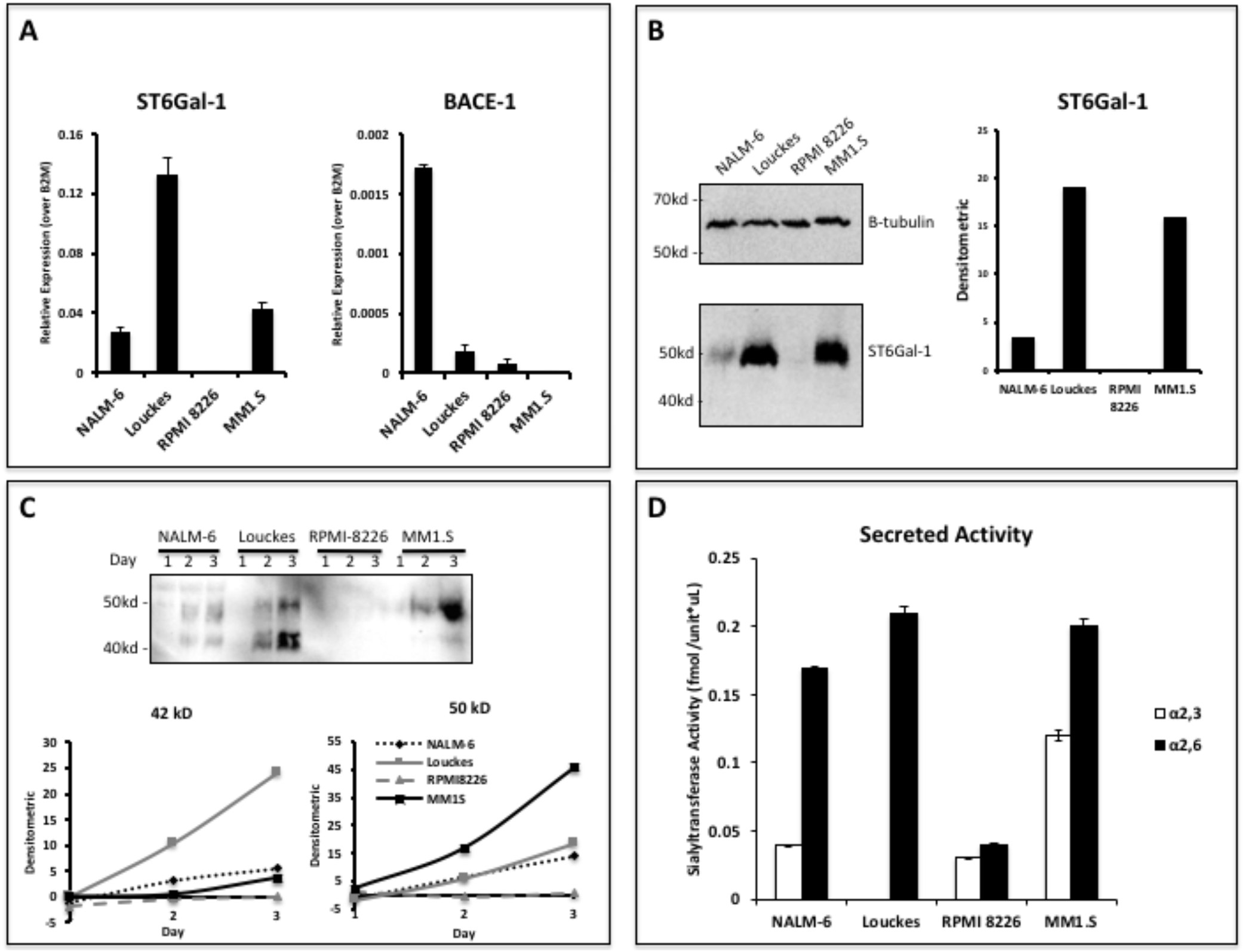
Human B lymphoblastoid cells secrete ST6Gal-1 of 50kD and 42kD size. Human lymphoblastoid cell lines derived from the pre-B (NALM-6), germinal center (Louckes) and plasma cell (RPMI 8226 and MM1.S) stages were profiled for ST6Gal-1 expression and secretion. (A) RT-qPCR analysis of ST6Gal-1 and beta-secretase BACE-1 mRNA (n=3 technical replciates) (B) Intracellular ST6Gal-1 protein levels, analyzed by western blot (left) and quantified (right). (C) Protein levels of ST6Gal-1 in the serum-free conditioned medium of cell cultures 1-3 days after plating 10^6 cells/ml, analyzed by western blot (top) and quantified for 50 kD and 42 kD sizes (bottom). (D) Sialyltransferase activity in conditioned medium, relative to media only control, as determined by incorporation of ^3H^NeuAc into Gal(β4)GlcNAc-O-Bn acceptor. The α2,6-sialyl product was quantified by SNA-agarose column chromatography. Data is representative of multiple experiments with similar results.

Theoretically, extracellular ST6Gal-1 can enzymatically reconstruct sialic acid on cell surfaces in the presence of a sialic acid donor substrate [36]. In order to determine if B cell secreted ST6Gal-1 is capable of extrinsically remodeling cell surfaces, we applied concentrated conditioned medium from Louckes cells (concentrated ∼50X) to metabolically fixed, sialidase pretreated human hepatoma HepG2 cells, in the presence or absence of the sialic acid donor substrate CMP-sialic acid for 1hr. The conditioned medium alone did not restore SNA reactivity, but required the presence of the sugar donor substrate to result in a striking increase in cell surface SNA reactivity (*Fig* 2). These results indicate that the sialyltransferase secreted by B cells is enzymatically active and capable of extrinsic sialylation of cell surface glycans.

**Figure 2.**
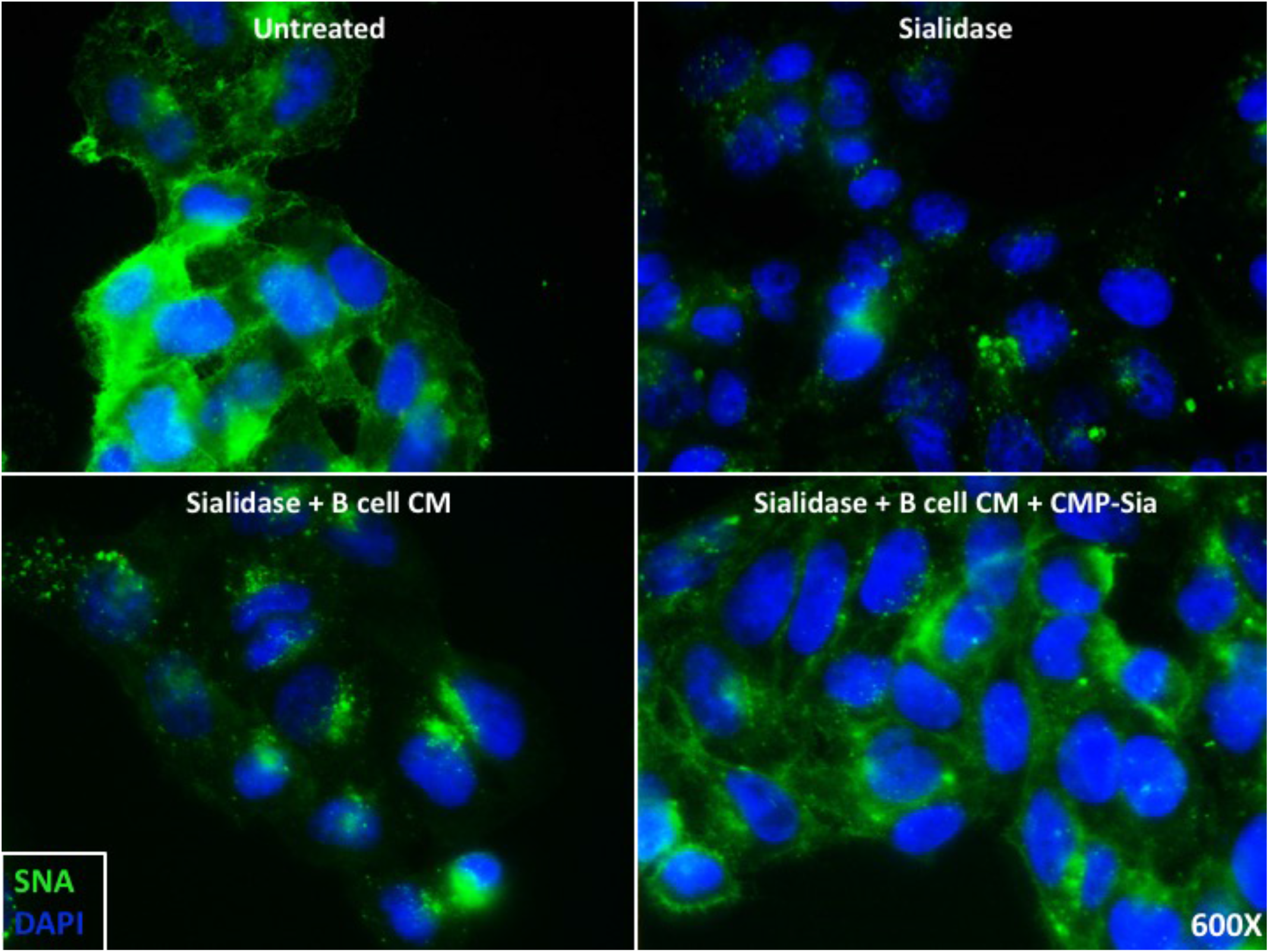
B cell conditioned medium extrinsically restores SNA reactivity of target cells. HepG2 human liver cells were grown on glass cover slides, fixed (10 min in 5% formalin) to disable endogenous metabolism. Cells were treated with *C. perfringens* sialidase C (Roche) for 1 hour at 37C to remove cell surface sialic acid, then incubated with concentrated (∼50X) B cell conditioned medium (CM) from Louckes grown in serum-free medium, in the presence or absence of 0.05 mM CMP-sialic acid.

### B cells glycosylate c-kit+ hematopoietic progenitors and suppress granulopoiesis in co-culture

Mice deficient in circulatory ST6Gal-1 exhibit exaggerated neutrophilia inducible by various inflammatory stimuli [29; 30; 37]. Supplementation of recombinant ST6Gal-1 is sufficient to blunt development of G-CSF and IL-5 dependent colonies from whole bone marrow cells [49]. This is due to sialylation of the multipotent granulocyte/monocyte progenitor (GMP), leading to reduced expression of myeloperoxidase, C/EBP-α, Gr-1, and neutrophil production [27]. Early B cells, mature B cells, and antibody-producing plasma cells represent a significant fraction of bone marrow cells. Therefore, we hypothesized that B cell-derived extrinsic ST6Gal-1 participates in the modification of neighboring hematopoietic stem and progenitor cells (HSPC) to suppress granulopoiesis.

To test this, human B lymphoblastoid cell lines were co-cultured with murine HSPCs (c-Kit+) and SCF, IL-3, TPO, Flt-3, G-CSF, and CMP-sialic acid for 3 days. In order to resolve the two populations by flow cytometry, the murine cells were labeled with the plasma membrane dye CellTrace Violet (*Fig 3a*). Co-cultures were generated at 1:1 and 4:1 ratio of B cells to HSPCs. Co-culture of B cells with HSPC resulted in modifications to cell surface sialic acid levels, as measured by reactivity towards the lectins *Sambucus nigra* (SNA) and *Maackia amurensis II* (MAL-II), which recognizes α2,6- and α2,3-sialic acids, respectively (*Fig 3b*). Furthermore, only ST6Gal-1 secreting B cell lines (NALM-6, Louckes, MM1.S) were able to increase cell-surface SNA reactivity of HSPC in a dose-dependent manner. In contrast, changes in MAL-II reactivity had no apparent correlation with ST6Gal-1 status, and only exhibited dose-dependent changes in RPMI 8226 co-cultures. We also assessed expression of CD11b and Gr-1, markers of myeloid differentiation, on the murine HSPC after co-culture (*Fig 3c*). After co-culture, we observed that ST6Gal-1 expressing B cells were able to modify expression of the granulocyte-specific marker Gr-1, but not the pan-myeloid marker CD11b on the murine HSPCs. When the frequencies of Gr-1+ cells from all co-cultures were plotted against SNA reactivity, there was a striking negative correlation (r^2^ = 0.66, p < 0.0001) (*Fig 3d*). This pattern was not present when frequencies of CD11b+ myeloid cells were plotted against SNA reactivity (r^2^ = 0.04, p = 0.2491). Based on these observations, we conclude that B cells are able to modify α2,6-sialylation and granulocytic differentiation of HSPCs in co-culture, and that the α2,6-sialylation of HSPCs was strongly and negatively associated with a key granulocytic marker.

**Figure 3.**
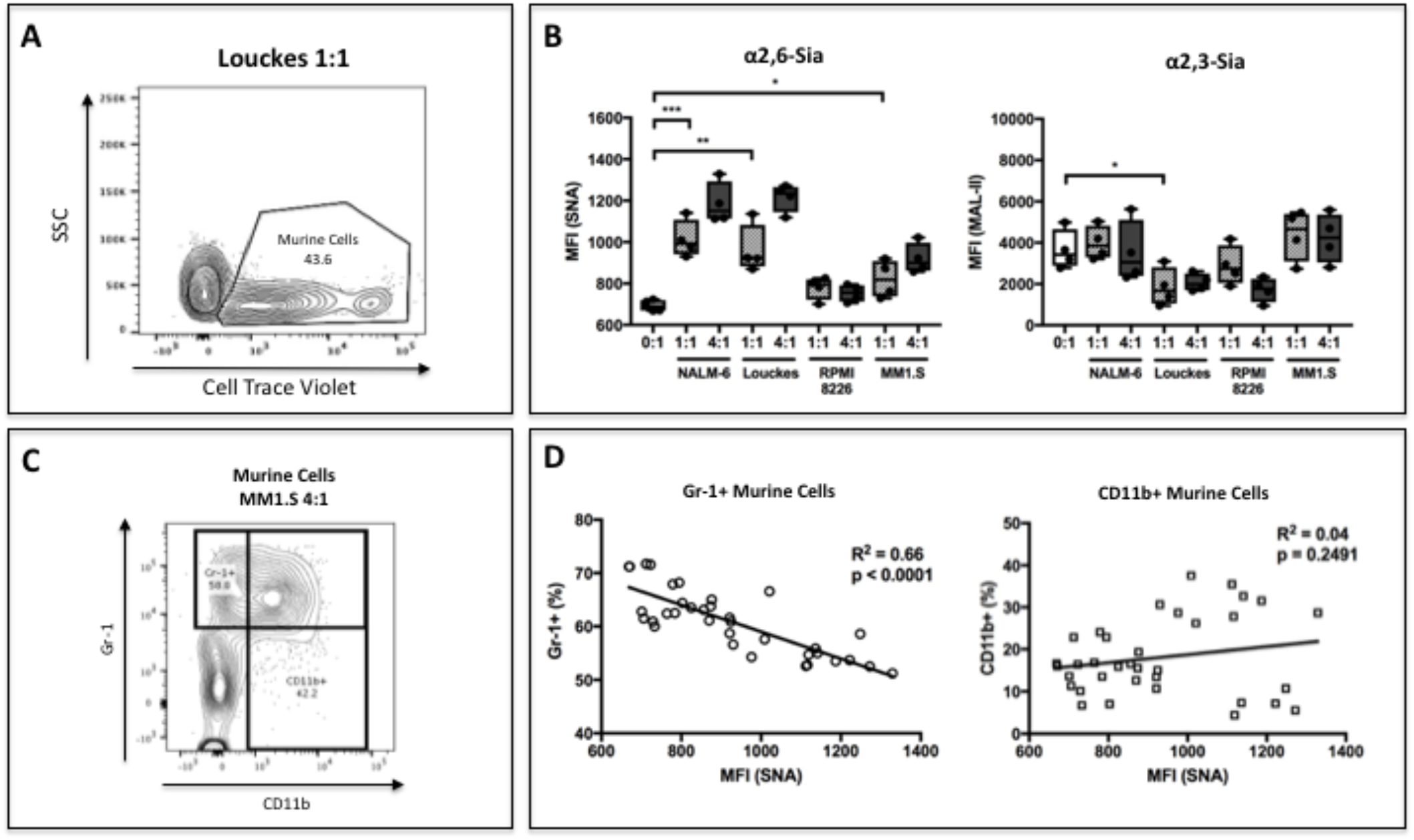
B cells modify HSPC SNA reactivity and Gr-1 expression in co-culture. Human B lymphoblastoid cell lines were co-cultured with CellTrace Violet-labeled murine c-kit+ bone marrow cells, at indicated ratios of 1:1 or 4:1, for 3 days with SCF, IL-3, G-CSF, TPO, and Flt-3 (A) Representative resolution of B cells and murine HSPCs by flow cytometry, with the HSPCs staining positive for CellTrace Violet. (B) Levels of SNA (α2,6-Sia) and MAL-II (α2,3-Sia) reactivity on the HSPCs in monoculture or co-culture with indicated B cells. (C) Flow cytometric separation of CD11b+ and Gr1+ cells from murine HSPCs, after 3 days of co-culture with human B lymphoblastoid cells. (D) Correlation between SNA reactivity and frequency of murine Gr-1+ or CD11b+ cells (expressed as % of total CellTrace Violet+ cells). Data is from a single experiment representative of three individual experiments, with n=4 technical replicates per condition; *p<0.05, **p<0.01, ***p<0.001 by student’s T-test.

### B cells secrete ST6Gal-1 to modify non-self hematopoietic cells *in vivo*

Our data demonstrate that human leukocytes in the B cell lineage can release functional ST6Gal-1 that modifies the glycosylation and granulopoietic differentiation of co-cultured hematopoietic progenitors *in vitro*. However, we wished to understand whether hematopoietic cells are a significant source of extracellular ST6Gal-1 *in vivo*. To answer this, we performed bone marrow transplantation of either wild type (C57BL/6) or *St6gal1*-KO whole bone marrow into *St6gal1*-KO /μMT recipients. These recipients were ST6Gal-1 deficient as well as B cell deficient due to the loss the heavy chain of IgM (Ighm-/-). In this scheme, all ST6Gal-1 present in extracellular compartments must originate from the donor hematopoietic cells (*Fig 4a*). Since the acquisition of IgM expression in the B cell lineage roughly coincides with the exit of immature B cells from the bone marrow prior to their development into naïve, recirculating mature B cells, all mature B cells recovered from the chimeras also must be of donor origin. In order to ascertain if ST6Gal-1 had been secreted, we measured sialyltransferase activities in the blood of the chimeras up to 8 weeks after transplantation using artificial O-benzyl-linked sialyltransferase substrates. Blood α2,3-sialyltransferase activity, which did not depend on ST6Gal-1, was not changed. In contrast, α2,6-sialyltransferase activity was entirely dependent upon ST6Gal-1 expression in the donor hematopoietic cells, and this was steadily increased with time (*Fig 4b*). Blood ST6Gal-1 activity even reached the baseline wild-type average by 8 weeks post-transplant, demonstrating that hematopoietic cells alone, without contribution from liver, were sufficient to maintain baseline blood ST6Gal-1.

**Figure 4.**
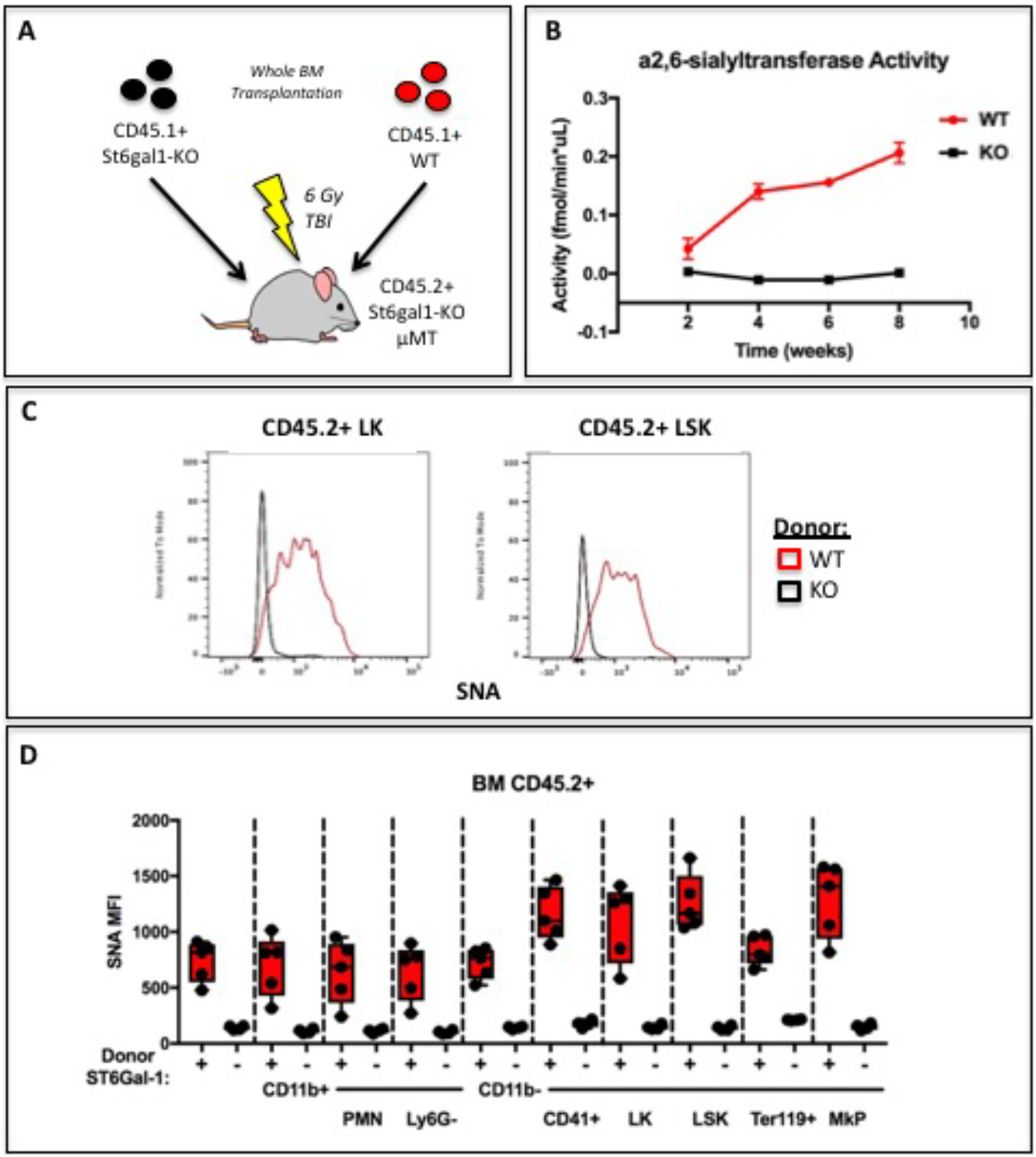
Hematopoietic Cells Supply Extracellular ST6Gal-1 for Extrinsic Sialylation *in vivo*. (A) CD45.1+ ST6Gal-1 sufficient (WT) or deficient (KO) whole bone marrow was used to reconstitute irradiated CD45.2+/μMT/St6gal1-KO mice. (B) α2,6-sialyltransferase activity was quantified in serum of bone marrow chimeras at indicated time points. (C) Representative histograms of SNA-reactivity are shown for Lin-/c-kit+/Sca-1- (LK) and Lin-/c-kit+/Sca-1+ (LSK) progenitor compartments in the bone marrow, 8 weeks post-transplant. (D) Mean fluorescence intensity of SNA in CD45.2+ recipient bone marrow cell subsets was quantified by flow cytometry (n=5). All cell types were significantly (p<0.01) different between WT and KO donor chimeras by student’s T-test. Data is from a single experiment and representative of two individual experiments.

In order to distinguish between donor and recipient hematopoietic cells, we utilized CD45.1+ mice as donors, and CD45.2+ / *St6gal1*-KO /μMT mice as recipients. Amongst the CD45.2+ residual host-derived CD11b+ myeloid, c-kit+ hematopoietic progenitors, and CD41+ megakaryocyte-lineage cells, all of them unable to express their own ST6Gal-1, increases in SNA reactivity were noted (*Fig 4c, 4d*). Particularly, progenitor Lin-/c-kit+/Sca-1- (LK) and Lin-/c-kit+/Sca-1+ (LSK) populations, as well as CD41+ megakaryocyte lineage progenitors, were dramatically modified by cell non-autonomous extrinsic ST6Gal-1, in comparison to CD11b+ myeloid cell subtypes and Ter119+ erythrocyte progenitors. These obeservations suggest target-dependent bias in the activity of extracellular ST6Gal-1.

In order to understand the spatial distribution of α2,6-sialic acid, we visualized SNA reactivity within the chimeric marrow of B cell deficient, *St6gal1*-KO/μMT recipients reconstituted with WT marrow cells. Formalin-fixed and frozen whole femurs were stained with FITC-conjugated SNA lectin. These chimeras, in which functional ST6Gal-1 and B lineage cells can only come from the donor, exhibited heterogenous areas of SNA reactivity across the bone marrow. This data suggested areas of preferred enzymatic sialylation in the bone marrow environment. Hence, areas representing varying SNA positivity, designated as Regions of Interest (ROIs) were selected for analysis. Four such ROIs are outlined in white, with observed SNA staining intensity of Region 1 > Region 4 > Region 3 > Region 2 (*Fig 5a, top*). The whole femur was further stained for IgD (mature recirculating B cells from donor) and PF4 (megakaryocytes) (*Fig5a, middle*). The co-localization of donor-derived IgD+ B cells and PF4+ megakaryocytes within the pre-selected ROIs was examined. ROIs that were SNA-bright tend to have both megakaryocytes and IgD+ B cells, whereas the ROIs with poor SNA staining were notably deficient in megakaryocytes, IgD+ B cells, or both. This is represented in magnified images of Region 1 and Region 2, representing the highest and lowest regions based on SNA-reactivity (*Fig 5a, bottom*). Numerous IgD+ and PF4+ cells were observed colocalizing in Region 1, but far fewer PF4+ cells in Region 2. Quantitative analysis of the ROIs is summarized in *Fig 5b*. The analysis demonstrate that the number of IgD+ B cells and PF4+ megakaryocytes are positively correlated with SNA-reactivity of ROIs, and that the two cell types also tended to colocalize with each other. Distinctly, SNA+ cells were not limited to the IgD+ B cells, but often encompassed groups of cells in their vicinity, consistent with their extrinsic modification. Collectively, this information suggests strongly that not only medullary B cells, but also megakaryocytes are participants in medullary extrinsic ST6Gal-1 sialylation. The contribution of megakaryocytes in medullary extrinsic sialylation is most intriguing, since megakaryocytes generate the platelets that supply activated sialic acid donor substrates for extramedullarly extrinsic sialylation [26; 36].

**Figure 5.**
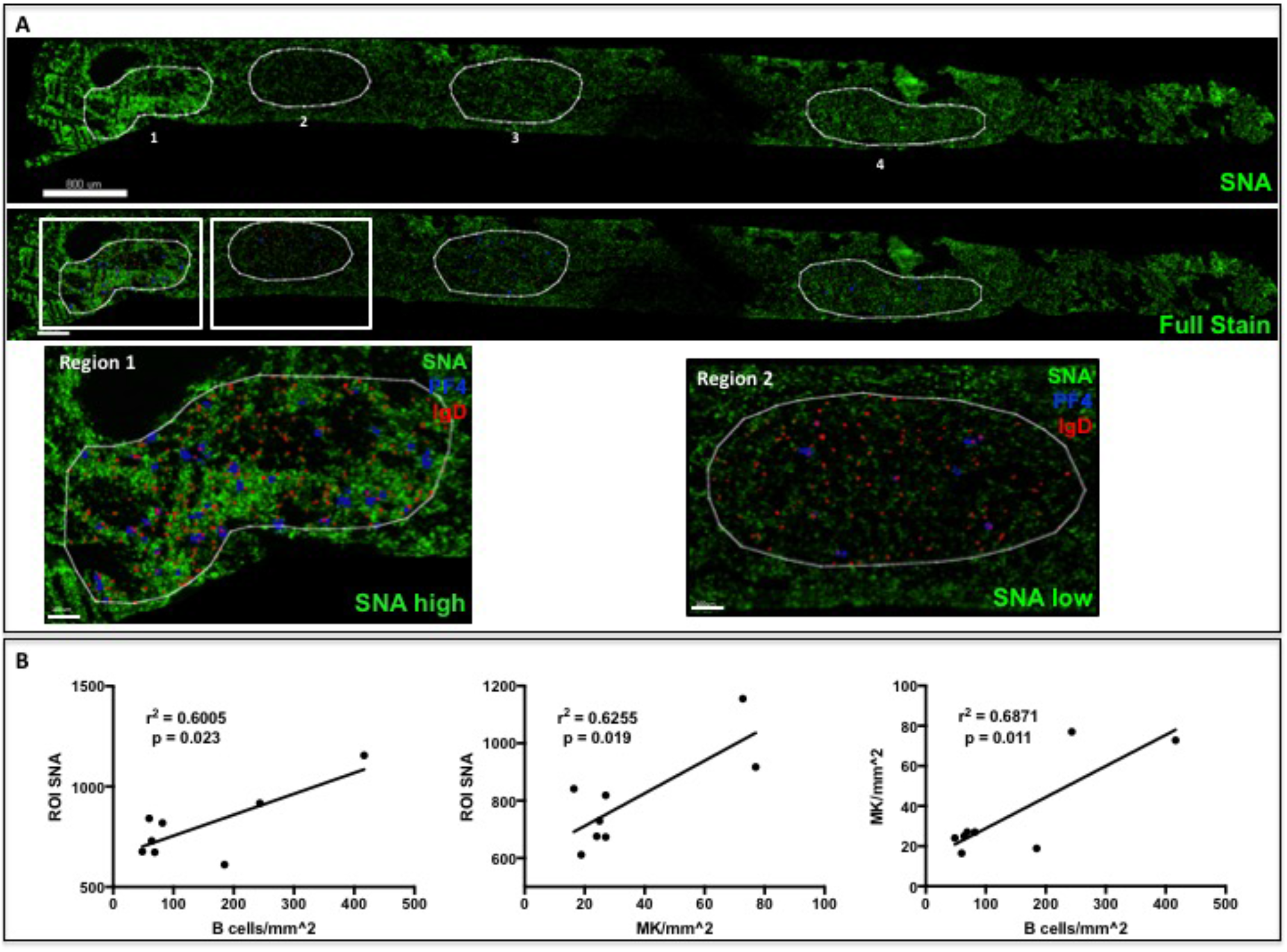
IgD+ B cells co-localize with Megakaryocytes to Regions of High a2,6-Sialylation within the Bone Marrow. (A) Wild-type bone marrow was allowed to reconstitute an irradiated μMT/ST6KO mouse for 8 weeks. Upon sacrifice, whole femurs were fixed, frozen, and sectioned for immunofluorescence staining. Heterogeneous SNA reactivity was observed and indicated regions of interest (ROI) were created with variable SNA reactivity (top panel). Megakaryocytes (PF4+) and mature B cells (IgD+) were identified within ROIs (middle panel and bottom panel insets). (B) Correlation of SNA reactivity of ROIs with IgD+ B cells (left panel) and PF4+ megakaryocytes (middle panel), as well as correlation between B cells and megakaryocytes within ROIs (right panel) from two chimeras. Data is derived from two biological replicates.

Though our observations demonstrate the feasibility for hematopoietic cells to drive extrinsic sialylation by ST6Gal-1, our histologic data strongly suggest B cells may be responsible for a portion of this effect. To test directly the idea that B lymphocytes can secrete a significant portion of ST6Gal-1 for extrinsic sialylation, CD45.2+ μMT recipients were irradiated and reconstituted with either syngeneic μMT bone marrow cells or an equal mix of μMT and Ighm+/+ wild-type bone marrow cells. The donor wild-type cells were CD45.1+, allowing them to be distinguished from the CD45.2+ μMT-derived cells during analysis (*Fig 6a*). In order to understand if the presence of Ighm+ B cells influenced the extracellular pool of ST6Gal-1, we analyzed serum samples for α2,6-sialyltransferase activity. Our data indicate that from week 6 post-transplant, mice reconstituted with B cells exhibited higher blood ST6Gal-1 activity (*Fig 6b*). Furthermore, at 10 weeks post-transplant, mice receiving Ighm+/+ donor bone marrow demonstrated increased SNA reactivity on the Ighm-/- hematopoietic cells within the blood and bone marrow, indicating that IgM+ B cells contribute to the sialylation of non-self hematopoietic cells (*Fig 6c*). Importantly, this occurred even in target cells expressing endogenous ST6Gal-1, indicating that the extrinsic sialylation process likely occurs naturally in wild-type, ST6Gal-1 expressing animals.

**Figure 6.**
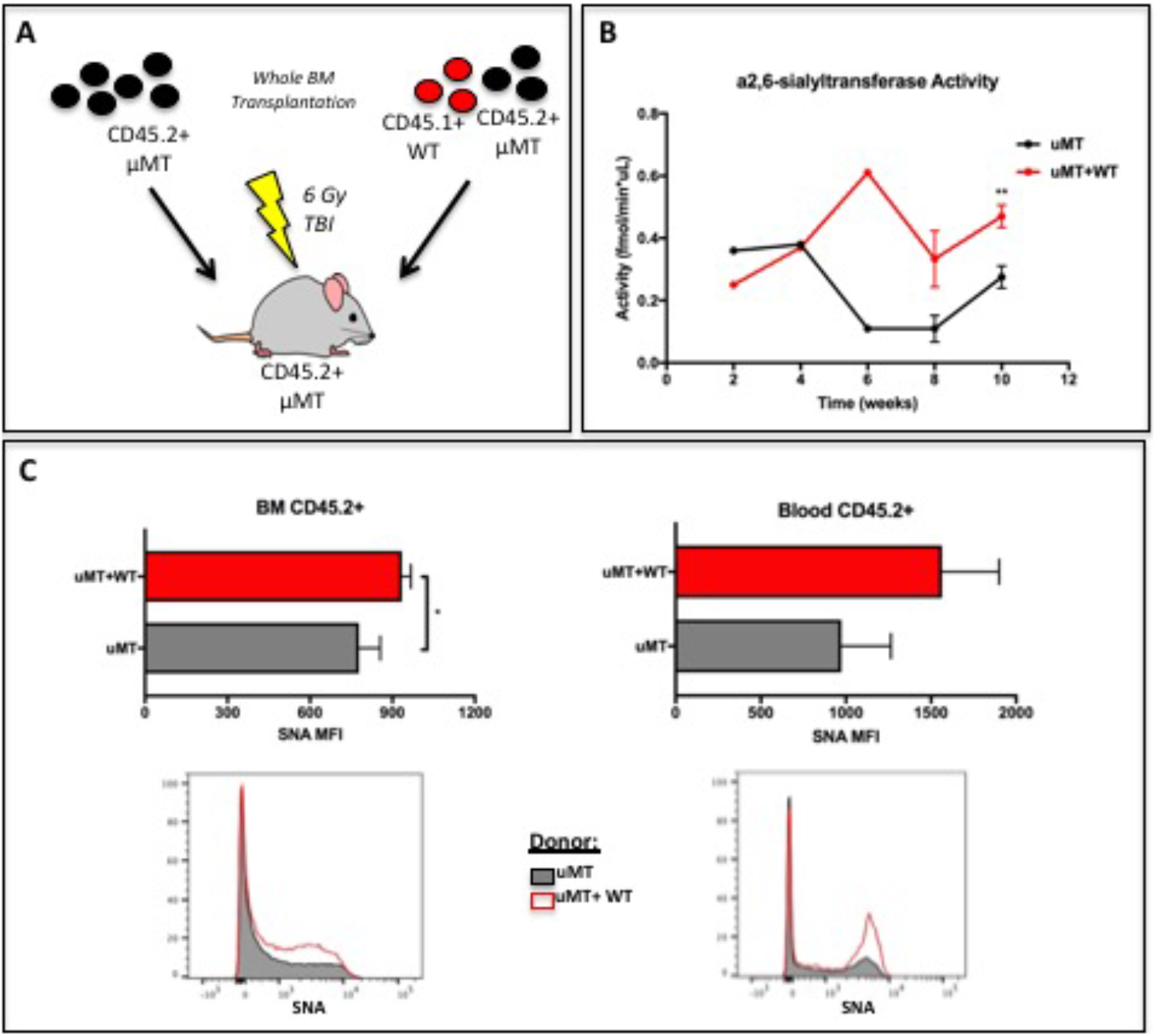
B cells Supply Extracellular ST6Gal-1 for Extrinsic Sialylation. (A) B cell-deficient μMT mice were irradiated and reconstituted with μMT (CD45.2+) or a mixture of μMT and WT (CD45.1+) whole BM. (B) a2,6-sialyltransferase activity in the serum was quantified over 10 weeks. (C) Bone marrow and peripheral blood CD45.2+ cells were analyzed for SNA reactivity at 10 weeks. Average SNA MFI and representative histograms of CD45.2+ μMT BM cells and peripheral blood cells are shown (grey= μMT donor only, red = μMT + WT donors). Data shown for one experiment with with n=3, but representative of two individual experiments. *p<0.05 **p<0.01 by student’s T-test.

### ST6Gal-1 expression in human multiple myeloma negatively correlates with bone marrow neutrophil abundance

B cells exist within the bone marrow medullary spaces at several stages during development. Whereas early B cells occupy the niche until successful arrangement of a functional BCR, mature B cells freely recirculate between the marrow, blood, and lymphoid tissues. In contrast, plasma cells can remain for years within the bone marrow to maintain systemic titers of protective antibodies, and are thought to occupy a specialized niche in proximity to megakaryocytes, eosinophils, and soluble pro-survival factors [50]. In order to understand if B cell-derived ST6Gal-1 is able to perturb normal HSPC development into mature leukocytes in a clinically relevant setting, we performed histological analyses of human bone marrow samples from freshly diagnosed, treatment-naïve patients with a plasma cell dyscrasia, multiple myeloma. Samples from only a limited number of patients (n=15) were available, and these were examined. Because of the clonal origin of multiple myeloma, the ST6Gal-1 expression of the neoplastic plasma cell could be assessed. Moreover, the high occupancy of the bone marrow by the neoplasm, which in some cases approached 60%, allowed for an assessment of the consequences of pathologically elevated ST6Gal-1 within the bone marrow microenvironment.

Paraffin-embedded bone marrow sections were stained for ST6Gal-1 using a DAB reagent. Plasma cell-specific expression of ST6Gal-1 was assessed according to intensity (1-5) and frequency (0-100%) by counting five groups of ten cells each in at least five fields of view per patient. The product of intensity and frequency is referred to as “ST6Gal-1 score”. ST6Gal-1 expression was highly heterogeneous between patients, and varied from nearly completely absent to intense expression in 100% of examined cells (*Fig 7a*). The expression of ST6Gal-1 in tumor cells was not associated with altered patient survival (*Fig 7b*). However, whereas the plasma cell burden and abundance of segmented neutrophils varied completely independently (r^2^ = 0.001, p = 0.88), we observed a striking relationship between plasma cell ST6Gal-1 expression and segmented neutrophils (*Fig 7c*). Qualitatively, patients with low ST6Gal-1 expression in plasma cells had evidence of abundant granulocytes on H&E staining, whereas high ST6Gal-1 expressing patients had far fewer visible granulocytes (*Fig 7d, arrows*). When ST6Gal-1 score was compared to the frequency of granulocyte lineage cells, as assessed by pathologist evaluation, we identified a strong negative correlation between ST6Gal-1 score and presence of segmented neutrophils (r^2^ = 0.42, p = 0.0083) (*Fig 7e*). Patients with low ST6Gal-1 scores (<100) had higher frequencies of mature neutrophils (22.32±4.11%), whereas those with high ST6Gal-1 scores (>100) had markedly lower frequency (9.1±2.6%) (*Fig 7e inset*).

**Figure 7.**
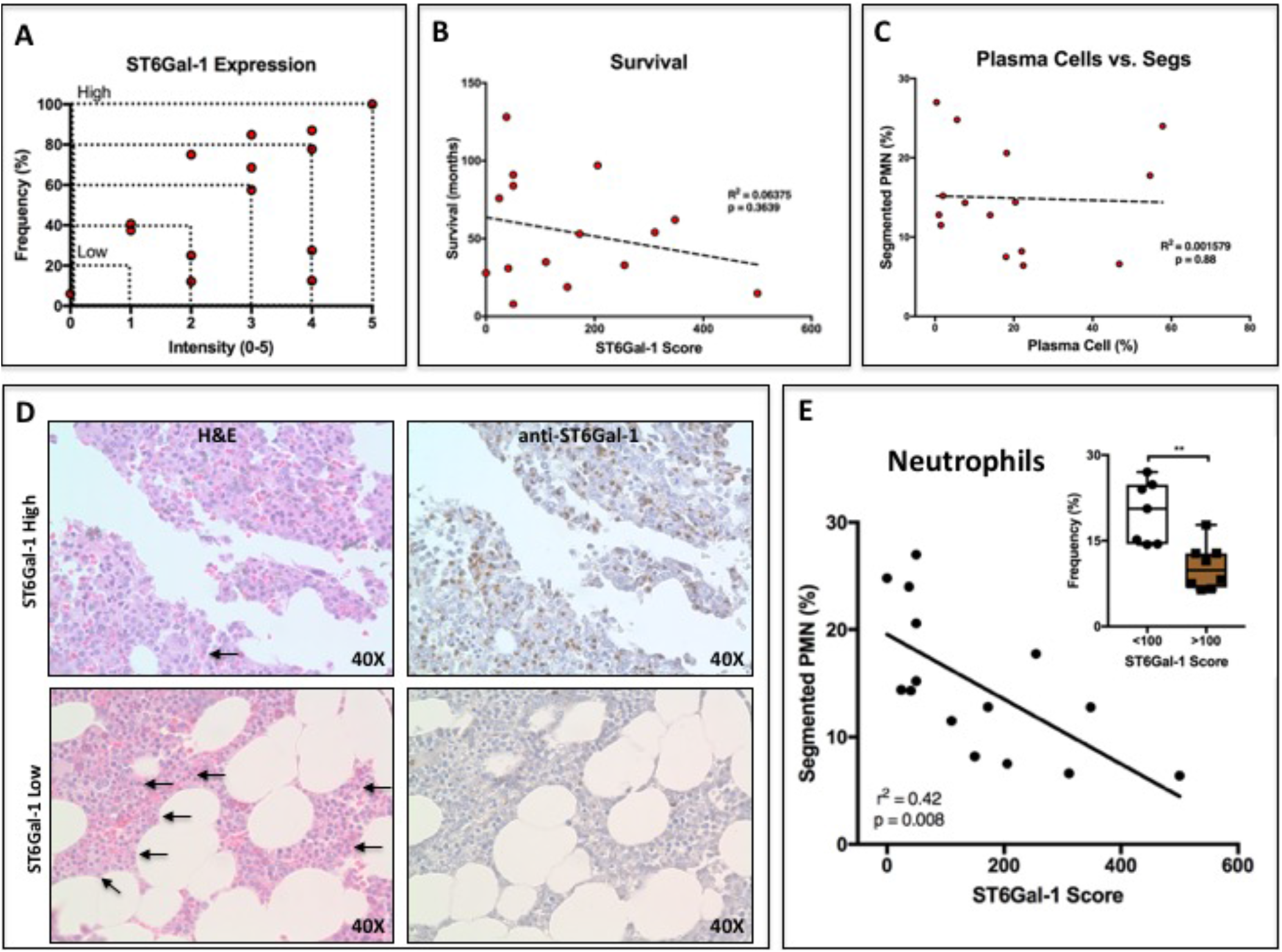
ST6Gal-1 Expression Levels in Human Multiple Myeloma Cells Correlates Negatively with Bone Marrow Neutrophil Abundance. (A) Quantification of ST6Gal-1 expression in bone marrow histological specimens from treatment-naive multiple myeloma patients (n=15). (B) Relationship between ST6Gal-1 expression and patient survival. (C) Relationship between Plasma cell abundance and segmented neutrophil abundance. (D) Representative data from one patient with high and one patient with low myeloma ST6Gal-1 expression, with accompanying H&E with neutrophils indicated (arrows). (E) Negative correlation between ST6Gal-1 expression and frequency of bands and segmented PMNs (p<0.01). Stratification of patients into low and high ST6Gal-1 expression was predictive of abundance of segmented neutrophils (**p<0.01, student’s T-test).

## Discussion

Effective cross-talk between components of the medullary environment is central to the demand-driven production of different lineages of blood cells. During systemic inflammation, B cells vacate the bone marrow due to the release of TNF-α and downregulation of CXCL12, salvaging space in preparation for the emergency generation of granulocytes [51]. However, it remains unclear if the departure of marrow B cells merely creates physical space for granulopoiesis or also disinhibits intrinsic granulopoietic processes. Our data support the existence of a paracrine signaling relationship, mediated by B cells via a novel extracellular glycosylation pathway, to influence the generation of granulocytes. This molecular pathway involves the release of catalytically active sialyltransferase ST6Gal-1 from B cells, and may contribute to the reciprocal inhibition of granulopoiesis during homeostasis. Our observations add to the body of literature documenting the emerging regulatory relationship between B cells and neutrophils. The newly-defined ‘B-helper’ subset of neutrophils (N_BH_) can prime marginal zone B cells for antibody production during inflammation by the release of BAFF [52; 53]. Conversely, evidence suggests that B cells inhibit neutrophil functions in a number of ways, including blocking chemotaxis and initiating β2 integrin-mediated apoptosis [54; 55]. B cell suppression of neutrophilic influx into the liver and spleen is necessary for the resolution of infections with a number of intracellular organisms, where neutrophils can cause damaging local and systemic inflammation [56; 57; 58]. Here, our observations suggest that B cells c influence the generation of new neutrophils by regulating glycosylation within the bone marrow, which are consistent with a separate body of literature documenting the development of excessive neutrophilic inflammation due to ST6Gal-1 insufficiency [27; 29; 30; 33; 49].

While localized gradients of cytokines, chemokines, and growth fators are already understood to create functional and developmental marrow niche spaces [59], our recent data also implicate a role for extracellular glycan-modifying enzymes such as ST6Gal-1 in the marrow hematopoietic environment. These include demonstrations of a profound role for extracellular, distally produced ST6Gal-1 in muting the transition of granulocyte-monocyte progenitors (GMP) to granulocyte progenitors (GP) [27], and that intravenously infused recombinant ST6Gal-1 can attenuate demand granulopoiesis in a mouse model of acute airway inflammation [31].

The existence of sialyltransferases within the extracellular milieau, particularly the blood, has been known for quite some time. Upregulation of serum ST6Gal-1 during inflammation was attributed to the induction of hepatic expression, leading to the designation of ST6Gal-1 as an acute phase reactant [17; 18; 39; 60; 61]. However, it was also recognized that in addition to hepatocytes, B cells are sophisticated expressers of ST6Gal-1, utilizing multiple tissue-specific transcripts during development [42; 46; 62]. Given the widespread distribution of ST6Gal-1 expressing mature B cells within secondary lymphoid tissues, blood, and bone marrow, we hypothesized that this population could be contributing to the extracellular pool of ST6Gal-1, thus regulating the sialylation and development of other hematopoietic cells. The observations in this study demonstrate that human B cells secrete functional ST6Gal-1 and are capable of modifying hematopoietic stem and progenitor cells (HSPC) in co-culture conditions to suppress granulocytic differentiation. *In vivo*, both hematopoietic and B cell-derived ST6Gal-1 are significant modifiers of bone marrow cell sialylation. Indeed, the limited endogenous expression of ST6Gal-1 in hematopoietic stem and progenitor populations may be compensated for by secreted, extracellular enzyme, making the bone marrow microenvironment a distinct niche space for extrinsic sialylation [25]. In contrast to untreated wild-type mice, we observed striking regional heterogeneity in bone marrow sialylation among *St6gal1*-KO chimeras reconstituted with wild-type bone marrow, with higher sialylation generally at the epiphysis and metaphysis of long bones. Interestingly, these sites of high sialylation contained high frequencies of IgD+ donor mature B cells that release the ST6Gal-1. We also observed spatial co-localization with megakaryocytes. In light of our previous observations that platelets can drive extrinsic sialylation in the periphery, it stands to reason that megakaryocytes, the ontogenic precursors to platelets [26], could provide the required sugar donor for this process in the bone marrow. Moreover, megakaryocytes emerge as organizers of the bone marrow niche, providing secreted HSC regulatory components, including TGF-β and CXCL4 [63; 64; 65]. It remains to be determined if megakaryocytes, like their peripheral ontogenic descendents [26], can release sialic acid donor substrates to influence medullary hematopoietic decisions.

ST6Gal-1 has been implicated in a variety of biological processes relevant to the development of disease, particularly in systemic inflammation [18] and in metastatic diseases [66]. The sialylation of IgG by ST6Gal-1 is necessary for the anti-inflammatory effects of IVIG therapy in autoimmune disease, and variations in serum IgG sialylation have been widely associated with inflammatory diseases [67; 68; 69]. Human GWAS studies have also associated genetic variation in ST6GAL1 with IgA nephropathy and flucloxacillin-induced liver damage [70; 71]. In epithelial carcinomas, ST6Gal-1 expression confers increased resistance to chemotherapy, hypoxia, and nutrient deprivation by promoting a stem-like phenotype, bolstering signaling through pro-survival and pro-proliferative EGFR, HIF-1α, and NF-κB pathways [72; 73; 74; 75; 76; 77]. Numerous early studies in both rodent models and humans have also documented concurrent increases in serum protein sialylation and sialyltransferase activity during malignancy, including in multiple myeloma, implying that ST6Gal-1 expressing tumor cells are capable of secreting into the extracellular pool [6; 78; 79; 80; 81; 82; 83]. In line with this, we observed that the hematopoietic compartment was able to completely restore the serum pool of ST6Gal-1 after bone marrow transplantation, and that B cells contributed a significant portion of this serum ST6Gal-1 activity. These results strongly argue in favor of a tissue-agnostic model of extrinsic sialylation, wherein extra-hepatic cell types secrete ST6Gal-1 into circulation. The functional consequences of fluctuations in blood ST6Gal-1 are yet unexplored, especially in malignancy. This current work examines specifically how cancer-derived ST6Gal-1 can perturb the generation of granulocytes, the best documented biologic role associated with extrinsic ST6Gal-1 sialylation [25; 30; 31; 49]. Multiple myeloma was examined because of the natural localization of tumor cells in the marrow, in proximity to nearby healthy HSPCs. ST6Gal-1 expression in MM varied dramatically from patient to patient. We observed that the abundance of segmented granulocytes within the marrow was strikingly and negatively associated with the level of ST6Gal-1 expression, but not with the overall abundance of multiple myeloma plasma cells. The data support the hypothesis that ST6Gal-1 negatively regulates granulocyte production in surrounding healthy progenitors, independently of disease burden.

Recent work in our group suggests that extrinsic ST6Gal-1 may have a broad ability to coordinate the development and function of multiple immune cell types [27; 28; 31]. Given the well-documented role of immune cells in cancer, further investigation into the role of extrinsic glycosylation in cancer is merited. The strong negative association between human multiple myeloma ST6Gal-1 expression and neutrophil prevalence indicates that tumor-derived ST6Gal-1 may dysregulate the development of bystander immune cells in the tumor microenvironment, with a variety of potential implications. At the very least, the ability of cancer-derived ST6Gal-1 to disrupt granulopoiesis predicts a diminished capacity for the patient to combat bacterial infections. Furthermore, the correlation between ST6Gal-1 levels and worse patient outcomes in a number of other cancers may be in part due to the extrinsic modification of mature tumor-associated leukocytes. This is consistent with reports that myeloid cell surface α2,6-sialylation diminishes maturation, activation, antigen cross-presentation and anti-tumor immune responses in dendritic cells, for example [84; 85; 86]. Collectively, our data hint at biologic effects of ST6Gal-1 in cancer that extend beyond the cell-intrinsic modulation of oncogenic signaling pathways, instead being mediated by a novel, extrinsic axis of glycosylation, and galvanized by the growing importance of immune cells in malignancy.

## Materials & Methods

### Animal models

The *St6gal1*-KO strain has been backcrossed 15 generations onto a C57BL/6J background and maintained at Roswell Park’s Laboratory Animal Shared Resource (LASR) facility. The B cell deficient B6.129S2 – Ighm^tm1Cgn^/J mouse μMT (The Jackson Laboratory) was used as a donor and recipient in bone marrow transplantation. The reference CD45.1 expressing wild-type strain used was B6.SJL-Ptprc^a^ Pepc^b^/BoyJ, in order to distinguish donor cells from recipient mice, which express the CD45.2 allele of the Ptprc locus. For transplantations, mice received 6 Gy whole body gamma-radiation and were rescued with 4.0×10^6^ whole bone marrow cells from a single donor or two donors equally. Mice were euthanized after 8-10 weeks for analysis. Unless otherwise indicated, mice between 7-10 weeks of age were used, and both sexes were equally represented. Roswell Park Institute of Animal Care and Use Committee approved maintenance of animals and all procedures used.

### Antibodies

For immunoblots and immunohistochemistry, anti-ST6Gal-1 (R&D Biosystems), anti-β-tubulin (Cell Signaling Technology), anti-PF4 (Peprotech), anti-IgD (eBioscience clone 11-26c), SNA-FITC (Vector labs) were used. For flow cytometry, SNA-FITC (Vector Labs), biotinylated MAL-II (Vector Labs), anti-Gr1-PE/Cy5, anti-CD11b-BV711, anti-CD45.2-PE/Cy7, anti-CD45.1-PerCP/Cy5.5, anti-Ly6G-APC, anti-Ter119-BV510, anti-CD41-BV421, anti-c-kit-APC/Cy7, and anti-Sca-1-PE (all Biolegend) were used.

### Analysis of cell lines

Human B lymphoblastoid cell lines were grown in RPMI base medium supplemented with 10% non-mitogenic heat-inactivated fetal bovine serum. All analyses were performed during logarithmic growth phase, and cell lines were kept in passage for no more than 6 weeks.

For RNA analysis, cells were washed, pelleted, and resuspended in TRI Reagent (MRC Inc.) and RNA extracted according to manufacturer’s instructions. 1.0μg RNA was converted to cDNA (iSCRIPT kit, Bio-rad), and then amplified by qPCR (iTaq Universal SYBR Green, Bio-rad) with intron-spanning primers towards human ST6Gal-1 and BACE1. Relative expression (2^dCt^) was calculated in reference to B2-microglobulin. Primer sequences are as follows: B2M: F 5’-GTGCTCGCGCTACTCTCTCT–3’, R 5’-TCAATGTCGGATGGATGAAACCC–3’; ST6GAL1: F 5’-CCTTGGGAGCTATGGGACATTC–3’, R 5’-TATCCACCTGGTCACACAGC–3’; BACE1: F 5’-TCTTCTCCCTGCAGCTTTGT–3’, R 5’-CAGCGAGTGGTCGATACCT–3’.

For intracellular protein, cells were washed, pelleted, and resuspended in RIPA cell lysis buffer with protease inhibitors, and 5-10ug of total protein resolved on 10% SDS gels, transferred onto activated PVDF membranes, and blocked in 5% fat-free milk for 1 hour. Blots were probed with primary antibody overnight at 4C, then washed and incubated with HRP-conjugated secondary for 1 hour. Membranes were developed using Pierce ECL WB Substrate (Thermo Scientific) and imaged using ChemiDoc Touch (Bio-rad). For analysis of secreted protein, cells were seeded at a density of 10^6^ cells/ml in serum-free RPMI, and cell-free conditioned medium collected after 24, 48, and 72 hours. In order to control for secreted protein per cell, an equal volume (1%) of conditioned media was resolved by 10% SDS-PAGE. Densitometric quantification of adjusted band intensity was performed separately for 50kD and 42kD forms of ST6Gal-1 using ImageJ software.

Enzymatic activity within conditioned medium was quantified using an artificial O-benzyl conjugated Gal-β1,4-GlcNAc acceptor, as has been described before [36]. Briefly, conditioned medium was incubated at 37C with artificial acceptor molecule and tritium-labeled CMP-sialic acid for 1 hour. The resulting reaction mix was applied to an O-benzyl reactive column and extensively washed, then eluted with methanol. Radioactive counts in the sample were quantified using a Beckman Coulter LS 6500 scintillation counter. α2,6-sialylated product was precipitated with SNA-agarose beads, and SNA-reactive fraction once again counted to quantify α2,6-sialyltransferase activity. Remaining α2,3-sialyltransferase activity was inferred.

### Extrinsic sialylation of fixed hepatocytes

HepG2 cells (ATCC) were seeded onto sterile glass cover slips in 6-well dishes for 3 days. Wells were washed with PBS and fixed for 5 minutes in 5% formalin solution. Cover slips were carefully removed from wells, and subjected to 1 hour treatment with bacterial sialidase C (Roche) at 37C, followed by incubation with concentrated Louckes conditioned medium for 1-2 hours at 37C, in the presence or absence of 100μM CMP-sialic acid charged sugar donor (EMD Millipore). Cover slips were blocked in 5% BSA, stained with SNA-FITC lectin, further stained with DAPI, then mounted onto charged microscope slides in 10% glycerol. Fluorescence was visualized immediately using a Nikon Eclipse E600 microscope with EXFO X-cite 120 light source. Spot RT3 camera and Spot Software were used to capture images.

### LK (Lin^neg^ cKit^pos^) cell co-culture

St6gal1-KO mouse bone marrow mononuclear cells were obtained and enriched for c-Kit+ cells using MACS columns (Miltenyi Biotechnology). Resulting Lin-neg:cKit+ (LK) hematopoietic progenitors (HSPCs) were stained for 20 minutes at room temperature with CellTrace Violet (Thermo Fisher), as per manufacturer’s instructions. Stained cells were quenched with media and washed thoroughly before quantification, and 10,000 cells cultured in 96-well round-bottom plates with either 10,000 or 40,000 human B lymphoblastoid cells at logarithmic growth phase. To induce differentiation and proliferation, cultures were supplemented with recombinant SCF (50ng/ml; BioVision), G-CSF (20ng/ml; Peprotech), IL-3 (5ng/ml; BioVision), TPO (25ng/ml; Peprotech), and FLT-3 (30ng/ml; Peprotech). After three days, cells were analyzed by flow cytometry for CellTrace Violet to discriminate between HSPCs and B cells, and murine cells further analyzed for cell surface glycans and expression of granulocyte markers. Flow cytometry data was acquired with BD LSR II flow cytometer and analyzed with FlowJo software.

### Analysis of bone marrow chimeras

Femurs of indicated bone marrow transplantation chimeras were flushed extensively to obtain cells. Peripheral blood was collected from the retro-orbital venous plexus in citrate-containing anticoagulant. All samples were subjected to ACK lysis in order to remove anucleated cells, then stained with the appropriate combination of antibodies for 20 minutes, washed, and analyzed by BD LSR II flow cytometer. Data was analyzed with FlowJo software, and donor status of individual cells was distinguished by CD45.1 and CD45.2 staining.

### Histological analysis of whole murine femurs

Femurs were fixed in a paraformaldehyde–lysine–periodate fixative overnight (0.01 M Sodium-M-Periodate, 0.075M L-Lysine, 1% PFA), rehydrated in 30% sucrose in a phosphate buffer solution for 48 hours, embedded in OCT (TissueTek, Sakura), and snap frozen in an isopentane/dry ice mixture [87]. Whole longitudinal sections (7 μm) sections were obtained using a Leica Cryostat and the Cryojane tape transfer system. Tissue sections were thawed, rehydrated and permeabilized in Tris-buffered saline with 0.1% Tween (T-TBS), blocked with 5% BSA, then incubated with FITC-conjugated SNA lectin (VectorLabs) for 1h, then washed prior to incubation with the appropriate primary antibodies. These were followed by the corresponding Alexa Fluor secondary antibodies (1:500, Invitrogen). Fluorescence whole slide imaging was performed on a Nikon Eclipse Ti2. Quantification of SNA density staining, megakaryocytes and IgD+ cell localization and numeration analysis was executed using Imaris (Bitplane) and Matlab (MathWorks) software. SNA density staining of the marked regions of interest (ROIs) was normalized from different femurs by accounting for total SNA intensity in the whole bone scan.

### Histological analysis of human bone marrow

All experiments involving human samples were evaluated and approved by the Institutional Review Board (IRB) prior to their initiation. Banked human biospecimens were provided by the Disease Bank and BioRepository at Roswell Park Comprehensive Cancer Center under protocol BDR 082017. Human multiple myeloma samples, collected from newly diagnosed and treatment-naïve patients, were examined. Samples from 15 treatment-naïve patients were available from the Repository, and these were de-identified prior to transfer, and associated patient information and pathology reports were provided via an honest broker. Paraffin-embedded sections of human bone marrow were melted at 55C for 1 hour, twice dehydrated in xylene-containing HistoClear (National Diagnostics), then rehydrated in successive ethanol solutions, and heated in Antigen Unmasking Solution (Vector Labs) for 30 minutes. Slides were blocked in 5% BSA for 1 hour, incubated overnight with anti-ST6Gal-1 antibody, then with anti-goat-HRP secondary (R&D Biosystems) for 1 hour. Tissues were then immersed in Impact DAB stain (Vector Labs) for 120 seconds and rinsed in water for 3 minutes. Slides were counterstained for 60 seconds with hematoxylin. ST6Gal-1 expression was quantified by counting of positively-staining plasma cells and evaluation of intensity of staining in at least 5 fields of view per specimen, typically containing 50 cells each. Bone marrow plasma cells and neutrophils were quantified by pathologist evaluation at time of diagnosis and provided by the Pathology Shared Resource Network (PSRN).

### Statistical analyses

Experiments were conducted with a minimum sample size calculated for appropriate power to detect changes of at least 2-fold (α = 0.05, β = 0.80, SD = 0.5). Raw data is presented in all figures as mean±SD.

## Author Contributions

**EEI**: conceptualization, methodology, investigation, formal analysis, writing – original draft preparation, writing – review & editing. **MML-S** and **KMH**: conceptualization, methodology, formal analysis, resources, writing – review & editing, funding acquisition. **JTYL**: conceptualization, resources, writing – original draft preparation, writing – review & editing, supervision, project administration, funding acquisition.

## Acknowledgements

This work was supported by grant R01AI140736 (to JTYL), R01HL089224 (to KMH), K12HL141954 (to KMH and JTYL). The core facilities of Roswell Park Comprehensive Cancer Center used in this work were supported in part by NIH National Cancer Institute Cancer Center Support Grant CA076056. Additional support includes BRI Director’s Fellowship Award to MML-S. We would like to thank Jon Wieser for the computational analysis of bone marrow imaging.

## References

1. Y. Lee, M. Decker, H. Lee, and L. Ding, Extrinsic regulation of hematopoietic stem cells in development, homeostasis and diseases. Wiley Interdiscip Rev Dev Biol 6 (2017).

2. R. Chovatiya, and R. Medzhitov, Stress, inflammation, and defense of homeostasis. Mol Cell 54 (2014) 281–8.

3. L.M. Calvi, and D.C. Link, The hematopoietic stem cell niche in homeostasis and disease. Blood 126 (2015) 2443–51.

4. D.I. Gabrilovich, Myeloid-Derived Suppressor Cells. Cancer Immunol Res 5 (2017) 3–8.

5. Y.S. Kim, J. Perfomo, A. Bella, and J. Nordberg, Properties of a CMP-N-acetylneuraminic acid: glycoprotein sialyltransferase in human serum and erythrocyte membranes. Biochimica et Biophysica Acta 244 (1971) 505.

6. R.J. Bernacki, and U. Kim, Concomitant elevations in serum sialyltransferase activity and sialic acid content in rats with metastasizing mammary tumors. Science 195 (1977) 577.

7. C. Ip, Changes in tissue and serum sialyltransferase activities as related to proliferation and involution of the rat mammary gland. Biochimica et Biophysica Acta 628 (1980) 249.

8. M.M. Lee-Sundlov, D.J. Ashline, A.J. Hanneman, R. Grozovsky, V.N. Reinhold, K.M. Hoffmeister, and J.T. Lau, Circulating blood and platelets supply glycosyltransferases that enable extrinsic extracellular glycosylation. Glycobiology 27 (2017) 188–198.

9. C. Ip, and T. Dao, Alterations in serum glycosyltransferases and 5’-nucleotidase in breast cancer patients. Cancer Res 38 (1978) 723–8.

10. I.M. Evans, R. Hilf, M. Murphy, and H.B. Bosmann, Correlation of seurm, tumor, and liver serum glycoprotein: N-acetylneuraminic acid transferase activity with growth of the R3230AC mammary tumor in rats and relationship of the serum activity. Cancer Research 40 (1980) 3103.

11. M.M. Weiser, and J.R. Wilson, Serum levels of glycosyltransferases and related glycoproteins as indicators of cancer: biological and clinical implications. Critical Reviews in Clinical Laboratory Sciences 14 (1981) 189–239.

12. P.G. Berge, A. Wilhelm, H. Schriewer, and G. Wust, Serum-sialyltransferase activity in cancer patients. Klin Wochenschr 60 (1982) 445–9.

13. T.L. Dao, C. Ip, J.K. Patel, and G.S. Kishore, Serum and tumor sialyltransferase activities in women with breast cancer. Progress in Clinical & Biological Research 204 (1986) 31.

14. A.M. Cohen, D. Allalouf, M. Djaldetti, K. Weigl, N. Lehrer, and H. Levinsky, Sialyltransferase activity in plasma cells of multiple myeloma. Eur J Haematol 43 (1989) 191–4.

15. A. Magalhaes, H.O. Duarte, and C.A. Reis, Aberrant Glycosylation in Cancer: A Novel Molecular Mechanism Controlling Metastasis. Cancer Cell 31 (2017) 733–735.

16. J.G. Rodrigues, M. Balmana, J.A. Macedo, J. Pocas, A. Fernandes, J.C.M. de-Freitas-Junior, S.S. Pinho, J. Gomes, A. Magalhaes, C. Gomes, S. Mereiter, and C.A. Reis, Glycosylation in cancer: Selected roles in tumour progression, immune modulation and metastasis. Cell Immunol (2018).

17. H.A. Kaplan, B.M.R.N.J. Woloski, M. Hellman, and J.C. Jamieson, Studies on the effect of inflammation on rat liver and serum sialyltransferase: Evidence that inflammation causes release of Gal beta1-4 GlcNAc alpha2-6 sialyltransferase from liver. Journal of Biological Chemistry 258 (1983) 11505.

18. J.C. Jamieson, G. McCaffrey, and P.G. Harder, Sialyltransferase: a novel acute-phase reactant Comparative Biochemistry & Physiology - B: Comparative Biochemistry 105 (1993) 29.

19. E.V. Gracheva, N.K. Golovanova, M.V. Ezhov, P.P. Malyshev, and V.V. Kukharchuk, Plasma sialyltransferase activity in healthy subjects and atherosclerotic patients. Biochemistry (Moscow) 64 (1999) 1315.

20. T.M. Maguire, A.M. Gillian, D. O’Mahony, C.M. Coughlan, A. Dennihan, Breen, and Kc, A decrease in serum sialyltransferase levels in Alzheimer’s disease. Neurobiology of Aging 15 (1994) 99.

21. N. Malagolini, F. Dall’Olio, F. Serafini-Cessi, and C. Cessi, Effect of acute and chronic ethanol administration on rat liver alpha 2,6-sialyltransferase activity responsible for sialylation of serum transferrin. Alcoholism 13 (1989) 649.

22. M. Garige, M. Gong, and M.R. Lakshman, Ethanol destabilizes liver Gal beta l, 4GlcNAc alpha2,6-sialyltransferase, mRNA by depleting a 3’-untranslated region-specific binding protein. J Pharmacol Exp Ther 318 (2006) 1076–82.

23. M. Gong, M. Garige, K. Hirsch, and M.R. Lakshman, Liver Galbeta1,4GlcNAc alpha2,6-sialyltransferase is down-regulated in human alcoholics: possible cause for the appearance of asialoconjugates. Metabolism 56 (2007) 1241–7.

24. M. Gong, L. Castillo, R.S. Redman, M. Garige, K. Hirsch, M. Azuine, R.L. Amdur, D. Seth, P.S. Haber, and M.R. Lakshman, Down-regulation of liver Galbeta1, 4GlcNAc alpha2, 6-sialyltransferase gene by ethanol significantly correlates with alcoholic steatosis in humans. Metabolism 57 (2008) 1663–8.

25. M. Nasirikenari, L. Veillon, C.C. Collins, P. Azadi, and J.T. Lau, Remodeling of marrow hematopoietic stem and progenitor cells by non-self ST6Gal-1 sialyltransferase. J Biol Chem 289 (2014) 7178–89.

26. C.T. Manhardt, P.R. Punch, C.W.L. Dougher, and J.T.Y. Lau, Extrinsic sialylation is dynamically regulated by systemic triggers in vivo. J Biol Chem 292 (2017) 13514–13520.

27. C.W.L. Dougher, A. Buffone, Jr., M.J. Nemeth, M. Nasirikenari, E.E. Irons, P.N. Bogner, and J.T.Y. Lau, The blood-borne sialyltransferase ST6Gal-1 is a negative systemic regulator of granulopoiesis. J Leukoc Biol 102 (2017) 507–516.

28. E.E. Irons, and J.T.Y. Lau, Systemic ST6Gal-1 Is a Pro-survival Factor for Murine Transitional B Cells. Front Immunol 9 (2018) 2150.

29. M. Nasirikenari, E.V. Chandrasekaran, K.L. Matta, B.H. Segal, P.N. Bogner, A.A. Lugade, Y. Thanavala, J.J. Lee, and J.T. Lau, Altered eosinophil profile in mice with ST6Gal-1 deficiency: an additional role for ST6Gal-1 generated by the P1 promoter in regulating allergic inflammation. J Leukoc Biol 87 (2010) 457–66.

30. M. Nasirikenari, B.H. Segal, J.R. Ostberg, A. Urbasic, and J.T. Lau, Altered granulopoietic profile and exaggerated acute neutrophilic inflammation in mice with targeted deficiency in the sialyltransferase ST6Gal I. Blood 108 (2006) 3397–405.

31. M. Nasirikenari, A.A. Lugade, S. Neelamegham, Z. Gao, K.W. Moremen, P.N. Bogner, Y. Thanavala, and J.T.Y. Lau, Recombinant Sialyltransferase Infusion Mitigates Infection-Driven Acute Lung Inflammation. Front Immunol 10 (2019) 48.

32. M.B. Jones, D.M. Oswald, S. Joshi, S.W. Whiteheart, R. Orlando, and B.A. Cobb, B-cell-independent sialylation of IgG. Proc Natl Acad Sci U S A 113 (2016) 7207–12.

33. M.B. Jones, M. Nasirikenari, A.A. Lugade, Y. Thanavala, and J.T. Lau, Anti-inflammatory IgG production requires functional P1 promoter in beta-galactoside alpha2,6-sialyltransferase 1 (ST6Gal-1) gene. J Biol Chem 287 (2012) 15365–70.

34. J.D. Pagan, M. Kitaoka, and R.M. Anthony, Engineered Sialylation of Pathogenic Antibodies In Vivo Attenuates Autoimmune Disease. Cell (2017).

35. H.H. Wandall, V. Rumjantseva, A.L. Sorensen, S. Patel-Hett, E.C. Josefsson, E.P. Bennett, J.E. Italiano, Jr., H. Clausen, J.H. Hartwig, and K.M. Hoffmeister, The origin and function of platelet glycosyltransferases. Blood 120 (2012) 626–35.

36. M.M. Lee, M. Nasirikenari, C.T. Manhardt, D.J. Ashline, A.J. Hanneman, V.N. Reinhold, and J.T. Lau, Platelets support extracellular sialylation by supplying the sugar donor substrate. J Biol Chem 289 (2014) 8742–8.

37. M.M. Appenheimer, R.Y. Huang, E.V. Chandrasekaran, M. Dalziel, Y.P. Hu, P.D. Soloway, S.A. Wuensch, K.L. Matta, and J.T. Lau, Biologic contribution of P1 promoter-mediated expression of ST6Gal I sialyltransferase. Glycobiology 13 (2003) 591–600.

38. G. Lammers, and J.C. Jamieson, Studies on the effect of experimental inflammation on sialyltransferase in the mouse and guinea pig. Comparative Biochemistry & Physiology - B: Comparative Biochemistry 84 (1986) 181–7.

39. J.C. Jamieson, G. Lammers, R. Janzen, and B.M.R.N.J. Woloski, The acute phase response to inflammation: The role of monokines in changes in liver glycoproteins and enzymes of glycoprotein metabolism. Comparative Biochemistry and Physiology 87b (1987) 11–5.

40. X.C. Wang, T.J. Smith, and J.T. Lau, Transcriptional regulation of the liver beta-galactoside alpha 2,6-sialyltransferase by glucocorticoids. J Biol Chem 265 (1990) 17849–53.

41. M. Dalziel, S. Lemaire, J. Ewing, L. Kobayashi, and J.T. Lau, Hepatic acute phase induction of murine beta-galactoside alpha 2,6 sialyltransferase (ST6Gal I) is IL-6 dependent and mediated by elevation of exon H-containing class of transcripts. Glycobiology 9 (1999) 1003–8.

42. S.A. Wuensch, R.Y. Huang, J. Ewing, X. Liang, and J.T. Lau, Murine B cell differentiation is accompanied by programmed expression of multiple novel beta-galactoside alpha2, 6-sialyltransferase mRNA forms. Glycobiology 10 (2000) 67–75.

43. J. Muller, and L. Nitschke, The role of CD22 and Siglec-G in B-cell tolerance and autoimmune disease. Nat Rev Rheumatol 10 (2014) 422–8.

44. T. Hennet, D. Chui, J.C. Paulson, and J.D. Marth, Immune regulation by the ST6Gal sialyltransferase. Proc Natl Acad Sci U S A 95 (1998) 4504–9.

45. M. Dalziel, R.Y. Huang, F. Dall’Olio, J.R. Morris, J. Taylor-Papadimitriou, and J.T. Lau, Mouse ST6Gal sialyltransferase gene expression during mammary gland lactation. Glycobiology 11 (2001) 407–12.

46. X. Wang, A. Vertino, R.L. Eddy, M.G. Byers, S.N. Jani-Sait, T.B. Shows, and J.T. Lau, Chromosome mapping and organization of the human beta-galactoside alpha 2,6-sialyltransferase gene. Differential and cell-type specific usage of upstream exon sequences in B-lymphoblastoid cells. J Biol Chem 268 (1993) 4355–61.

47. S. Kitazume, Y. Tachida, R. Oka, K. Shirotani, T.C. Saido, and Y. Hashimoto, Alzheimer’s beta-secretase, beta-site amyloid precursor protein-cleaving enzyme, is responsible for cleavage secretion of a Golgi-resident sialyltransferase. Proc Natl Acad Sci U S A 98 (2001) 13554–9.

48. X. Deng, J. Zhang, Y. Liu, L. Chen, and C. Yu, TNF-alpha regulates the proteolytic degradation of ST6Gal-1 and endothelial cell-cell junctions through upregulating expression of BACE1. Sci Rep 7 (2017) 40256.

49. M.B. Jones, M. Nasirikenari, L. Feng, M.T. Migliore, K.S. Choi, L. Kazim, and J.T. Lau, Role for hepatic and circulatory ST6Gal-1 sialyltransferase in regulating myelopoiesis. J Biol Chem 285 (2010) 25009–17.

50. J.R. Wilmore, and D. Allman, Here, There, and Anywhere? Arguments for and against the Physical Plasma Cell Survival Niche. J Immunol 199 (2017) 839–845.

51. Y. Ueda, K. Yang, S.J. Foster, M. Kondo, and G. Kelsoe, Inflammation controls B lymphopoiesis by regulating chemokine CXCL12 expression. J Exp Med 199 (2004) 47–58.

52. P. Scapini, B. Nardelli, G. Nadali, F. Calzetti, G. Pizzolo, C. Montecucco, and M.A. Cassatella, G-CSF-stimulated neutrophils are a prominent source of functional BLyS. J Exp Med 197 (2003) 297–302.

53. I. Puga, M. Cols, C.M. Barra, B. He, L. Cassis, M. Gentile, L. Comerma, A. Chorny, M. Shan, W. Xu, G. Magri, D.M. Knowles, W. Tam, A. Chiu, J.B. Bussel, S. Serrano, J.A. Lorente, B. Bellosillo, J. Lloreta, N. Juanpere, F. Alameda, T. Baro, C.D. de Heredia, N. Toran, A. Catala, M. Torrebadell, C. Fortuny, V. Cusi, C. Carreras, G.A. Diaz, J.M. Blander, C.M. Farber, G. Silvestri, C. Cunningham-Rundles, M. Calvillo, C. Dufour, L.D. Notarangelo, V. Lougaris, A. Plebani, J.L. Casanova, S.C. Ganal, A. Diefenbach, J.I. Arostegui, M. Juan, J. Yague, N. Mahlaoui, J. Donadieu, K. Chen, and A. Cerutti, B cell-helper neutrophils stimulate the diversification and production of immunoglobulin in the marginal zone of the spleen. Nat Immunol 13 (2011) 170–80.

54. T.K. Kondratieva, E.I. Rubakova, I.A. Linge, V.V. Evstifeev, K.B. Majorov, and A.S. Apt, B cells delay neutrophil migration toward the site of stimulus: tardiness critical for effective bacillus Calmette-Guerin vaccination against tuberculosis infection in mice. J Immunol 184 (2010) 1227–34.

55. J.H. Kim, J. Podstawka, Y. Lou, L. Li, E.K.S. Lee, M. Divangahi, B. Petri, F.R. Jirik, M.M. Kelly, and B.G. Yipp, Aged polymorphonuclear leukocytes cause fibrotic interstitial lung disease in the absence of regulation by B cells. Nat Immunol 19 (2018) 192–201.

56. C.M. Bosio, and K.L. Elkins, Susceptibility to secondary Francisella tularensis live vaccine strain infection in B-cell-deficient mice is associated with neutrophilia but not with defects in specific T-cell-mediated immunity. Infect Immun 69 (2001) 194–203.

57. S.C. Smelt, S.E. Cotterell, C.R. Engwerda, and P.M. Kaye, B cell-deficient mice are highly resistant to Leishmania donovani infection, but develop neutrophil-mediated tissue pathology. J Immunol 164 (2000) 3681–8.

58. A.J. Buendia, L. Del Rio, N. Ortega, J. Sanchez, M.C. Gallego, M.R. Caro, J.A. Navarro, F. Cuello, and J. Salinas, B-cell-deficient mice show an exacerbated inflammatory response in a model of Chlamydophila abortus infection. Infect Immun 70 (2002) 6911–8.

59. A. Birbrair, and P.S. Frenette, Niche heterogeneity in the bone marrow. Ann N Y Acad Sci 1370 (2016) 82–96.

60. J.C. Jamieson, Studies on the effect of 1-deoxynojirimycin on the release of albumin, sialyltransferase and alpha 1 acid glycoprotein from liver slices from normal and inflamed rats. Life Sciences 43 (1988) 691.

61. W. van Dijk, W. Boers, M. Sala, A. Lasthuis, and S. Mookerjea, Activity and secretion of sialyltransferase in primary monolayer cultures of rat hepatocytes cultured with and without dexamethasone. Biochem.Cell.Biol. 64 (1986) 79.

62. N.W. Lo, and J.T. Lau, Novel heterogeneity exists in the 5’-untranslated region of the beta-galactoside alpha 2,6-sialytransferase mRNAs in the human B-lymphoblastoid cell line, louckes. Biochem Biophys Res Commun 228 (1996) 380–5.

63. I. Bruns, D. Lucas, S. Pinho, J. Ahmed, M.P. Lambert, Y. Kunisaki, C. Scheiermann, L. Schiff, M. Poncz, A. Bergman, and P.S. Frenette, Megakaryocytes regulate hematopoietic stem cell quiescence through CXCL4 secretion. Nat Med 20 (2014) 1315–20.

64. Y. Gong, M. Zhao, W. Yang, A. Gao, X. Yin, L. Hu, X. Wang, J. Xu, S. Hao, T. Cheng, and H. Cheng, Megakaryocyte-derived excessive transforming growth factor beta1 inhibits proliferation of normal hematopoietic stem cells in acute myeloid leukemia. Exp Hematol 60 (2018) 40–46 e2.

65. S. Pinho, and P.S. Frenette, Haematopoietic stem cell activity and interactions with the niche. Nat Rev Mol Cell Biol (2019).

66. J. Lu, and J. Gu, Significance of beta-Galactoside alpha2,6 Sialyltranferase 1 in Cancers. Molecules 20 (2015) 7509–27.

67. J.D. Pagan, M. Kitaoka, and R.M. Anthony, Engineered Sialylation of Pathogenic Antibodies In Vivo Attenuates Autoimmune Disease. Cell 172 (2018) 564–577 e13.

68. F. Nimmerjahn, and J.V. Ravetch, Anti-inflammatory actions of intravenous immunoglobulin. Annu Rev Immunol 26 (2008) 513–33.

69. M.H. Biermann, G. Griffante, M.J. Podolska, S. Boeltz, J. Sturmer, L.E. Munoz, R. Bilyy, and M. Herrmann, Sweet but dangerous - the role of immunoglobulin G glycosylation in autoimmunity and inflammation. Lupus 25 (2016) 934–42.

70. M. Li, J.N. Foo, J.Q. Wang, H.Q. Low, X.Q. Tang, K.Y. Toh, P.R. Yin, C.C. Khor, Y.F. Goh, I.D. Irwan, R.C. Xu, A.K. Andiappan, J.X. Bei, O. Rotzschke, M.H. Chen, C.Y. Cheng, L.D. Sun, G.R. Jiang, T.Y. Wong, H.L. Lin, T. Aung, Y.H. Liao, S.M. Saw, K. Ye, R.P. Ebstein, Q.K. Chen, W. Shi, S.H. Chew, J. Chen, F.R. Zhang, S.P. Li, G. Xu, E.S. Tai, L. Wang, N. Chen, X.J. Zhang, Y.X. Zeng, H. Zhang, Z.H. Liu, X.Q. Yu, and J.J. Liu, Identification of new susceptibility loci for IgA nephropathy in Han Chinese. Nat Commun 6 (2015) 7270.

71. A.K. Daly, P.T. Donaldson, P. Bhatnagar, Y. Shen, I. Pe’er, A. Floratos, M.J. Daly, D.B. Goldstein, S. John, M.R. Nelson, J. Graham, B.K. Park, J.F. Dillon, W. Bernal, H.J. Cordell, M. Pirmohamed, G.P. Aithal, C.P. Day, D. Study, and S.A.E.C. International, HLA-B*5701 genotype is a major determinant of drug-induced liver injury due to flucloxacillin. Nat Genet 41 (2009) 816–9.

72. M.J. Schultz, A.T. Holdbrooks, A. Chakraborty, W.E. Grizzle, C.N. Landen, D.J. Buchsbaum, M.G. Conner, R.C. Arend, K.J. Yoon, C.A. Klug, D.C. Bullard, R.A. Kesterson, P.G. Oliver, A.K. O’Connor, B.K. Yoder, and S.L. Bellis, The Tumor-Associated Glycosyltransferase ST6Gal-I Regulates Stem Cell Transcription Factors and Confers a Cancer Stem Cell Phenotype. Cancer Res 76 (2016) 3978–88.

73. C.M. Britain, K.A. Dorsett, and S.L. Bellis, The Glycosyltransferase ST6Gal-I Protects Tumor Cells against Serum Growth Factor Withdrawal by Enhancing Survival Signaling and Proliferative Potential. J Biol Chem 292 (2017) 4663–4673.

74. A. Chakraborty, K.A. Dorsett, H.Q. Trummell, E.S. Yang, P.G. Oliver, J.A. Bonner, D.J. Buchsbaum, and S.L. Bellis, ST6Gal-I sialyltransferase promotes chemoresistance in pancreatic ductal adenocarcinoma by abrogating gemcitabine-mediated DNA damage. J Biol Chem 293 (2018) 984–994.

75. A.T. Holdbrooks, C.M. Britain, and S.L. Bellis, ST6Gal-I sialyltransferase promotes tumor necrosis factor (TNF)-mediated cancer cell survival via sialylation of the TNF receptor 1 (TNFR1) death receptor. J Biol Chem 293 (2018) 1610–1622.

76. C.M. Britain, A.T. Holdbrooks, J.C. Anderson, C.D. Willey, and S.L. Bellis, Sialylation of EGFR by the ST6Gal-I sialyltransferase promotes EGFR activation and resistance to gefitinib-mediated cell death. J Ovarian Res 11 (2018) 12.

77. R.B. Jones, K.A. Dorsett, A.B. Hjelmeland, and S.L. Bellis, The ST6Gal-I sialyltransferase protects tumor cells against hypoxia by enhancing HIF-1alpha signaling. J Biol Chem 293 (2018) 5659–5667.

78. A.M. Cohen, D. Allalouf, H. Bessler, M. Djaldetti, T. Malachi, and H. Levinsky, Sialyltransferase activity and sialic acid levels in multiple myeloma and monoclonal gammopathy. Eur J Haematol 42 (1989) 289–92.

79. A.M. Cohen, T. Wacks, B.B. Lurie, and H. Levinsky, Interferon-alpha and dexamethasone effect on AF10 myeloma cell line sialytransferase activity. Biochemical Medicine & Metabolic Biology 50 (1993) 9.

80. H. Chelibonova-Lorer, S. Ivanov, E. Gavazova, and M. Antonova, Characterization of sialyltransferases from serum of normal and hepatoma Mc-29 bearing chickens. International Journal of Biochemistry 18 (1986) 271.

81. K. Dairaku, T. Miyagi, and A. Wakui, Increase in serum sialyltransferase in tumor bearing rats: The origin and nature of the increased enzyme. Gann 74 (1983) 656.

82. P. Gessner, S. Riedl, A. Quentmaier, and W. Kemmner, Enhanced activity of CMP-neuAc:Gal beta 1-4GlcNAc:alpha 2,6-sialyltransferase in metastasizing human colorectal tumor tissue and serum of tumor patients. Cancer Letters 75 (1993) 143.

83. T.C. Poon, C.H. Chiu, P.B. Lai, T.S. Mok, B. Zee, A.T. Chan, J.J. Sung, and P.J. Johnson, Correlation and prognostic significance of beta-galactoside alpha-2,6-sialyltransferase and serum monosialylated alpha-fetoprotein in hepatocellular carcinoma. World J Gastroenterol 11 (2005) 6701–6.

84. M. Silva, Z. Silva, G. Marques, T. Ferro, M. Goncalves, M. Monteiro, S.J. van Vliet, E. Mohr, A.C. Lino, A.R. Fernandes, F.A. Lima, Y. van Kooyk, T. Matos, C.E. Tadokoro, and P.A. Videira, Sialic acid removal from dendritic cells improves antigen cross-presentation and boosts anti-tumor immune responses. Oncotarget 7 (2016) 41053–41066.

85. H.J. Crespo, J.T. Lau, and P.A. Videira, Dendritic cells: a spot on sialic Acid. Front Immunol 4 (2013) 491.

86. M.G. Cabral, Z. Silva, D. Ligeiro, E. Seixas, H. Crespo, M.A. Carrascal, M. Silva, A.R. Piteira, P. Paixao, J.T. Lau, and P.A. Videira, The phagocytic capacity and immunological potency of human dendritic cells is improved by alpha2,6-sialic acid deficiency. Immunology 138 (2013) 235–45.

87. C. Nombela-Arrieta, G. Pivarnik, B. Winkel, K.J. Canty, B. Harley, J.E. Mahoney, S.Y. Park, J. Lu, A. Protopopov, and L.E. Silberstein, Quantitative imaging of haematopoietic stem and progenitor cell localization and hypoxic status in the bone marrow microenvironment. Nat Cell Biol 15 (2013) 533–43.

